# Autoantibody subclass predominance is not driven by aberrant class switching or impaired B cell development

**DOI:** 10.1101/2023.06.30.546522

**Authors:** Laurent M. Paardekooper, Yvonne E. Fillié-Grijpma, Alita J. van der Sluijs-Gelling, Mihaela Zlei, Remco van Doorn, Maarten H. Vermeer, Manuela Paunovic, Maarten J. Titulaer, Silvère M. van der Maarel, Jacques J.M. van Dongen, Jan J. Verschuuren, Maartje G. Huijbers, the T2B consortium

## Abstract

A subset of autoimmune diseases is characterized by predominant pathogenic IgG4 autoantibodies (IgG4-AIDs). Why IgG4 predominates in these disorders is unknown. We hypothesized that dysregulated B cell maturation or aberrant class switching causes overrepresentation of IgG4^+^ B cells and plasma cells. Therefore, we compared the B cell compartment of patients with muscle-specific kinase (MuSK) myasthenia gravis (MG), pemphigus, leucine-rich glioma inactivated (LGI1) encephalitis and contactin-associated protein-like 2 (CASPR2) encephalitis (four IgG4-AIDs) to patients with acetylcholine receptor (AChR) MG, Lambert-Eaton myasthenic syndrome (LEMS) (two IgG1-3-AIDs) and age-matched healthy donors, using flow cytometry. B cell subset relative abundance at all maturation stages was normal, except for a, possibly treatment-related, reduction in immature and naïve CD5^+^ cells in IgG4-AIDs. IgG4^+^ B cell and plasma cell fractions were normal in IgG4-AID patients, however they had an (sub)class-independent 8-fold increase in circulating mature CD20^-^CD138^+^ plasma cells. No autoreactivity was found in this subset after sorting. In conclusion, patients with IgG4-AID do not show increased numbers of IgG4-expressing cells. These results argue against aberrant B cell development in these patients and rather suggest the autoantibody subclass predominance to be antigen-driven. The similarities between B cell subset numbers among these patients suggest that these IgG4-AIDs, despite displaying variable clinical phenotypes, share a similar underlying immune profile.

**Graphical abstract:** 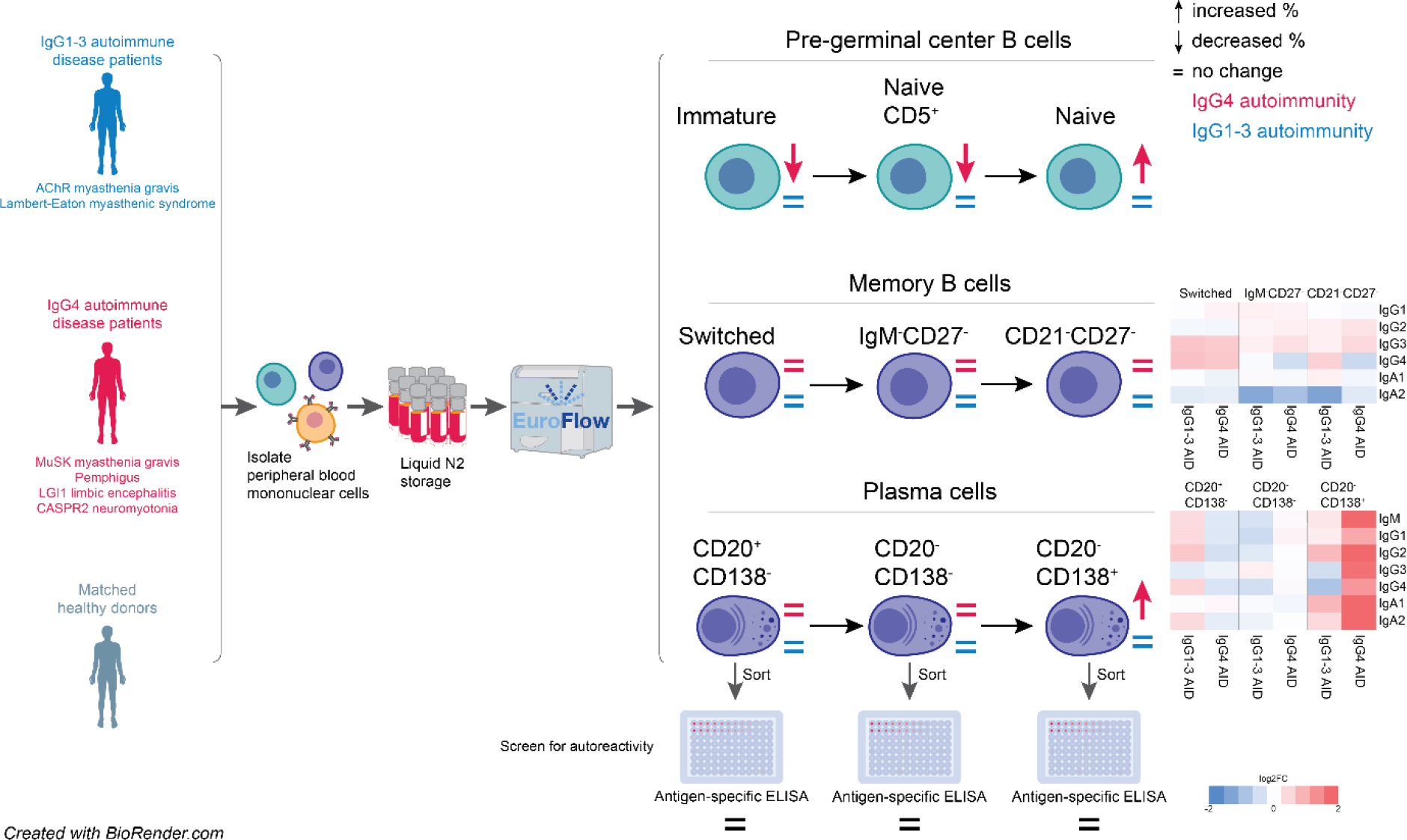

## 1. Introduction

An important factor that determines the pathophysiological mechanism in antibody-mediated autoimmune diseases is the dominant autoantibody (sub)class. The majority of antibody-mediated autoimmune diseases are caused by pro-inflammatory autoantibody subclasses such as ImmunoglobulinG 1 (IgG1) and IgG3 (1). These antibody subclasses, through activation of complement or immune cell-mediated cytotoxicity, damage the target organ causing the disease-associated symptoms (2). In addition, their bivalent nature allows them to crosslink their target antigens, often causing internalization and loss of surface antigen function which further contributes to the pathology (3, 4). Interestingly, in 2015 a group of autoimmune diseases (AIDs) predominated by autoantibodies of the IgG4 subclass was described (IgG4-AIDs) (5). This was surprising as IgG4 is generally considered an anti-inflammatory antibody subclass. IgG4 has low affinity for most Fc receptors and complement factor C1q (6–9). This means that IgG4 usually does not induce antibody-mediated phagocytosis, antibody-dependent cell-mediated cytotoxicity or complement-mediated tissue damage. Additionally, IgG4 antibodies are uniquely capable of Fab-arm exchange meaning exchange of antibody half molecules (one heavy chain and one light chain) resulting in bispecific, functionally monovalent IgG4 molecules (10–13). Because of the inability to activate the immune system and its relatively high affinity the effects of IgG4 are usually caused by blocking the function of the target antigen (5, 6, 14, 15).

To date, 29 different AIDs fit the criteria for IgG4-AID (16). These affect different organ systems and are generally rare with a prevalence of 0.001-5/10.000 individuals (17). During the last decades *in vitro* and *in vivo* studies have directly confirmed the pathogenicity of IgG4 autoantibodies in at least six IgG4-AIDs (18–21). Insight in the pathophysiology and immunological characteristics of these autoimmune diseases highlights several commonalities between these disorders: 1) IgG4 autoantibodies block essential protein-protein interactions thereby causing disease, 2) on a group level, IgG4 serum titers are only marginally increased (22–24), 3) they respond favorably to rituximab treatment (25–28) and 4) they show a strong association with HLA-class II haplotypes HLA-DQB1*05 and HLA-DRB1*04 (16, 29). These observations suggest that, although IgG4-AIDs affect different organs and cause a variety of symptoms, they may in fact share a similar underlying immunological profile.

Why IgG4 predominates in these autoimmune responses is poorly understood. This is relevant however as the switching to IgG4 may make autoantibodies more pathogenic (30) and treatment strategy may be adjusted accordingly. Class switching to IgG4 is known to occur in response to prolonged exposure to certain antigens such as bee venom (6, 31) or under influence of Th2 cytokines IL-4, IL-10 and IL-13 (32–35). Indeed, these cytokines were found increased in IgG4-AID patients and cross-reactivity was observed with autoantibodies from pemhigus patients with IgG4-inducing allergens (36–38). Lastly, dysregulated B cell maturation or aberrant class switching may cause overrepresentation of IgG4^+^ B cells and IgG4 plasma cells in immune responses. To further understand what is causing the IgG4 predominance in IgG4-AIDs, we investigated in detail the many IgH-isotype subsets of the circulating B cell compartment in four archetypical IgG4-AIDs and compared them to two IgG1-3-AIDs and age-matched healthy controls.

## 2. Results

### Study population

To investigate the role of abberant B cell development or class switching in subclass predominated AID, the B cell compartments of four IgG4-AIDs were immunophenotyped and compared to two IgG1-3-AIDs and healthy donors. An overview of the demographics of the study population is given in Table 1. Median age at time of blood draw and male:female ratio were comparable between groups. The CASPR2 encephalitis patient samples did not contain enough cells to perform a reliable in-depth phenotyping analysis (Sup. Fig. 3), therefore they were only included in the pre-germinal center analyses. One pemphigus patient sample was excluded due to an abnormally low total B cell count which can be attributed to the patient receiving rituximab infusion just before blood draw. In this study we aimed to include as many treatment naïve patients as possible or those only receiving low dosis of immunosuppression to limit a treatment bias. Several patients however received multiple treatments simultaneously. A complete description of the included study population is given in Sup. Table 1.

**Table 1.** Study population.

|  | Number of patients (excluded) | Age (median, min/max) | Sex (M:F) | Immunosuppressive treatment (number of patients) |
| --- | --- | --- | --- | --- |
| **MuSK Myasthenia Gravis** | 11 (0) | 59 (27-79) | 4:7 | None (2), prednisone (4), azathioprine (3), IVIG (1), cellcept (2), plasmaferesis (1), unknown (2) |
| **Pemphigus** | 10 (1) | 58 (26-75) | 5:4 | Prednisone (8), clobetasol (1) rituximab (1; excluded) |
| **LGI1 encephalitis** | 2 (0) | - (48-62) | 1:1 | Untreated (1), prednisone (1), IVIG (1) |
| **CASPR2 encephalitis** | 3 (included in pre-GC only) | 65 (57-69) | 3:0 | Untreated (1), azathioprine (2) |
| **Lambert-Eaton Myasthenic Syndrome** | 10 (0) | 56 (49 – 74) | 3:7 | None (7), prednisone (1), azathioprine (1), IVIG (1), hydrocortisone (1) |
| **AChR Myasthenia Gravis** | 10 (0) | 63 (18-79) | 7:3 | None (5), prednisone (5), IVIG (1) |
| **Healthy control** | 10 (0) | 58 (44-68) | 5:5 | - |
| **p-value (column statistic)** | - | 0.96 (1-way ANOVA) | 0.31 ($\chi^2$) | - |

### Decreased numbers of immature and naïve B cells in patients with AID may be treatment related

To investigate early B cell development stages we investigated the pre-germinal center (GC) B cells across all cohorts. The relative abundance of immature (Fig. 1A) and naïve CD5^+^ (Fig. 1B) B cells was lowered in MuSK MG and pemphigus patients compared to healthy controls. Consequently, their relative abundance of naïve CD5^-^ B cells is increased (Fig. 1C). LGI1 and CASPR2 encephalitis patients show a similar trend, but due to the low number of patients per group this analysis lacked power. Treatment status may influence immature B cell numbers (39). The observed reduction in cell numbers does not seem to be explained by the use of a single drug. Notably, the variance within the autoimmune disease groups was considerably larger than in healthy donors.

**Figure 1.**
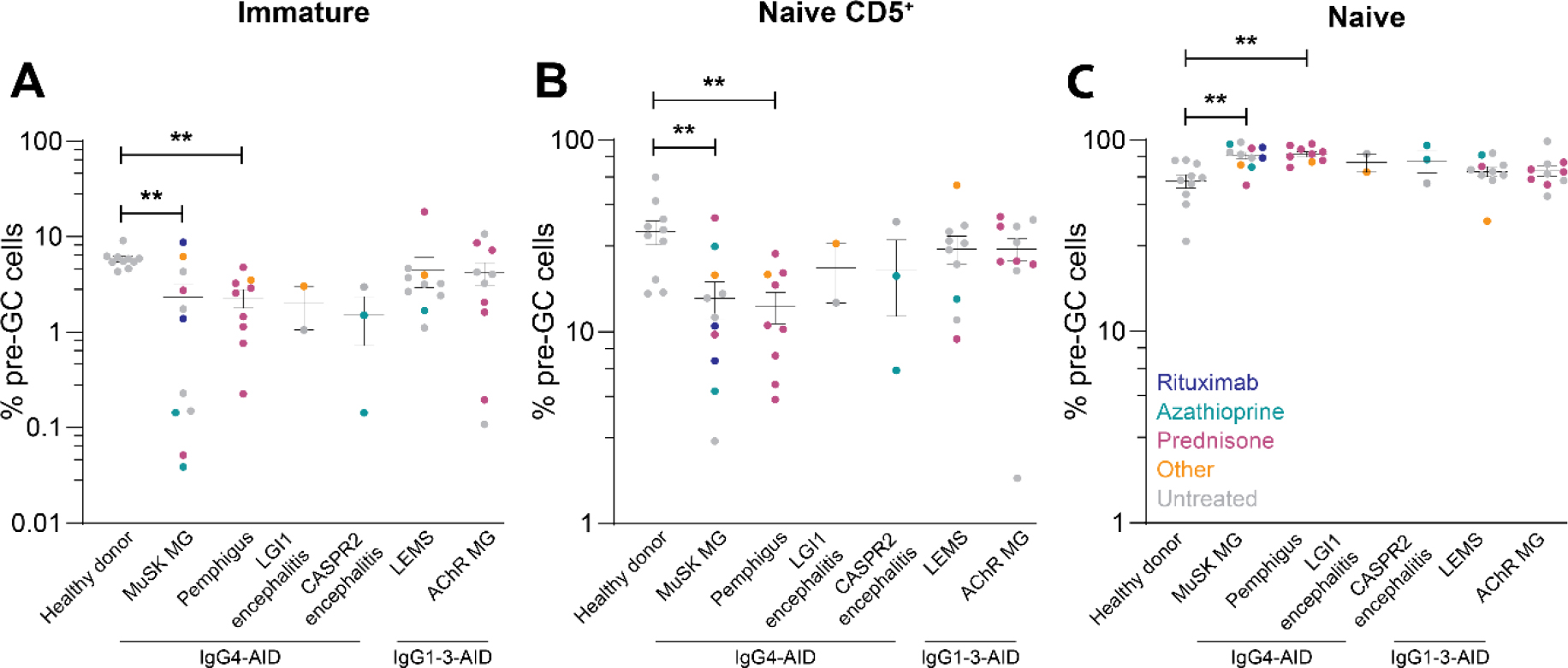
Pre-germinal center B cell fractions. **(A)** Immature, **(B)** naïve CD5^+^ and **(C)** naïve B cell counts as percentage of all pre-germinal center (GC) B cells. Treatment status is marked per patient. For all panels, IgG4-AIDs were compared to both healthy controls and IgG1-3-AIDs by one-way ANOVA followed by unpaired Student’s t-test (* p<0.05; ** p<0.01; *** p<0.005).

### B memory cell numbers are largely normal in IgG1-3-AID and IgG4-AID

GC formation is essential for the development of a functional antibody repertoire and is iniated by B cell receptor signaling after antigen encounter (40, 41). Pre-GC B cells are mostly of the IgM or IgD isotype and have not undergone affinity maturation yet as both class switching and somatic hypermutation take place in the GC (41–43). In the context of autoimmunity, post-GC antigen-experienced mature memory B cells and plasma cells are particularly of interest as they may harbor the autoreactive cell subsets. Overall total memory B cell levels are normal in all autoimmunity groups (Fig. 2A). The total number of switched (IgM^-^/D^-^CD27^+^) (Fig. 2B), double-negative (IgM^-^/D^-^CD27^-^) (Fig. 2C) and atypical (IgM^-^/D^-^CD21^-^CD27^-^) (Fig. 2D) memory B cells are similar to healthy donors. After stratification for B cell receptor (sub)class, total memory B cell (Fig. 2E), switched (Fig. 2F) and double-negative (Fig. 2G) memory B cell fractions are still similar between groups. Only in MuSK MG patients did we observe significantly lower atypical IgG4 B cells (IgM^-^/D^-^CD21^-^CD27^-^) compared to healthy controls and IgG1-3 AID patients (Fig. 2H). Surprisingly, atypical IgG4 memory B cells were increased in the IgG1-3 AIDs LEMS and AChR MG compared to IgG4-AID patients.

**Figure 2.**
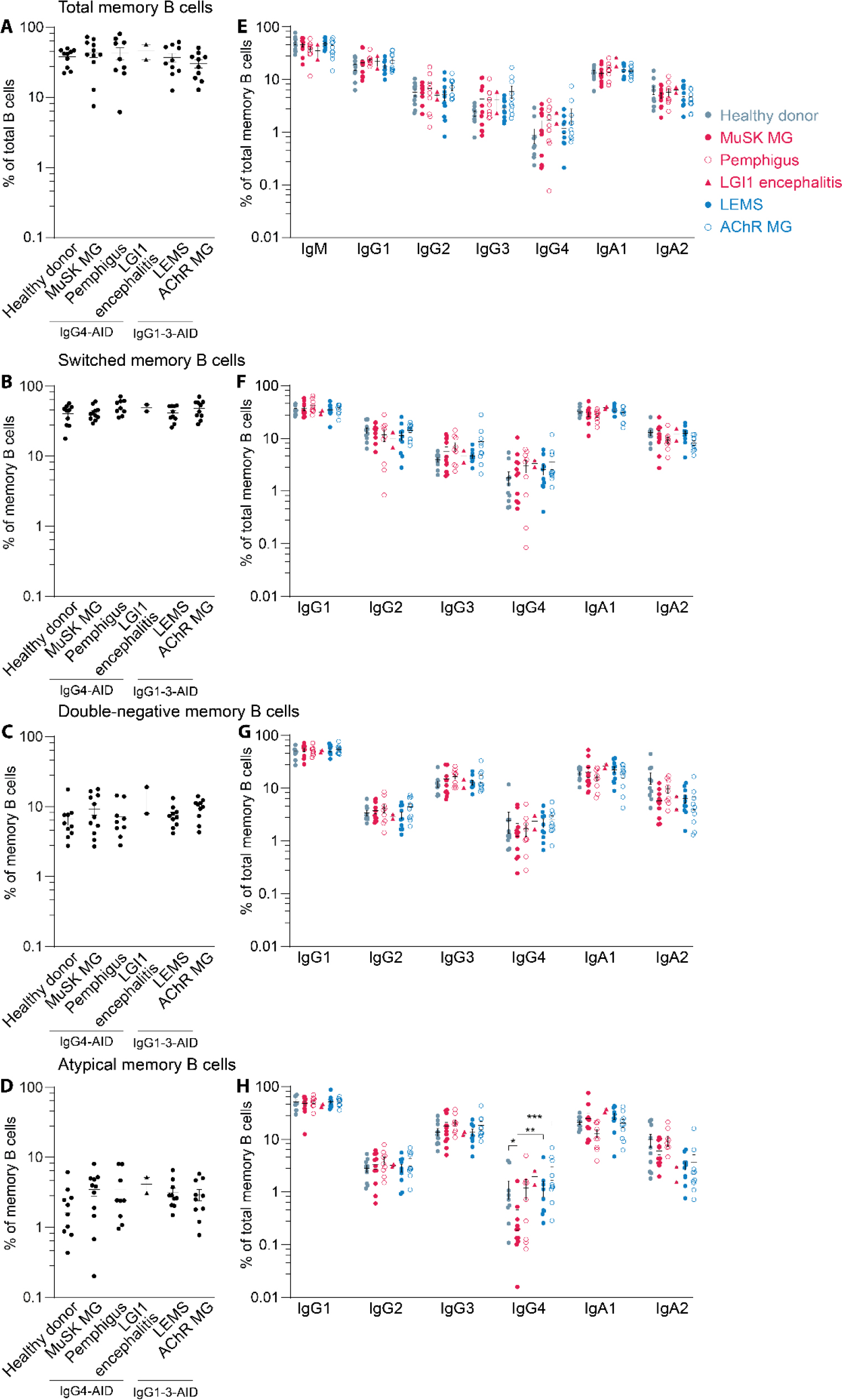
Memory B cell fractions of IgG4-AID and IgG1-3-AID patients. **(A)** Overall memory B cell (CD19^+^CD20^+^CD27^+^) counts normalized to total B cell counts, **(B)** switched memory B cell (IgM^-^/D^-^ CD19^+^CD20^+^CD27^+^) counts normalized to total B cell counts, **(C)** double-negative B cell (IgM^-^/D^-^ CD19^+^CD20^+^CD27^-^) counts normalized to total B cell counts, **(F)** atypical B cell (IgM^-^/D^-^ CD19^+^CD20^+^CD21^-^CD27^-^) counts normalized to total B cell counts. **(E), (F), (G), (H):** Respectively total memory, switched, double-negative and atypical B cell counts normalized to total memory B cell counts and stratified by B cell receptor (sub)class. For all panels, IgG4-AIDs were compared to both healthy controls and IgG1-3-AIDs by one-way ANOVA followed by unpaired Student’s t-test (* p<0.05; ** p<0.01; *** p<0.005).

### IgG4-AID patients have 8-fold increased circulating mature plasma cell numbers

Differentiating B cells can also commit to the plasma cell lineage upon leaving the GC (44). These plasmablasts express CD27 and CD38 as they fully mature into plasma cells they gradually lose expression of CD20 and gain expression of CD138 (45, 46). Total plasma cell fractions were comparable in all groups (Fig. 3A). When stratified by B cell receptor (sub)class, we observe increases in IgG1^+^ and IgG3^+^ plasma cells, as well as a decrease in IgA1^+^ plasma cells only seen in the pemphigus patients (Fig. 3B). Only IgG3^+^ plasma cells are increased in MuSK MG patients. There were no changes in IgG4^+^ plasma cell fractions for any of the IgG4-AIDs. Upon maturation to plasma cells, plasmablasts lose expression of CD20 while gaining expression of CD138 (47, 48). To investigate if IgG4-AIDs correlate with altered numbers of these matured plasma cells we quantified three specific plasma cell maturation stages: CD20^+^CD138^-^, CD20^-^CD138^-^ and CD20^-^CD138^+^. IgG1-3-AID patients show a slight reduction in CD20^-^CD138^-^ intermediate plasma cells (Fig. 4A). When stratified by B cell receptor (sub)class this reduction is observable in IgG1^+^ and IgG2^+^ plasma cells (Fig. 4B-C). At the same time, IgG1^+^ and IgG2^+^ CD20^+^CD138^-^ plasmablasts are increased in IgG1-3-AID patients. Interestingly, in all three IgG4-AID patient groups we observe increased fractions of the CD20^-^CD138^+^ fully matured plasma cells in comparison to both healthy controls and IgG1-3-AIDs (on average 8-fold increase, range 4-14; Fig. 4A).This increase is not specific to IgG4^+^ plasma cells and instead is observed in IgG1^+^ (pemphigus only), IgG2^+^, IgA1^+^ and IgA2^+^ (pemphigus only) CD20^-^CD138^+^ plasma cells (Fig. 4D, 4F, 4G, respectively).

**Figure 3.**
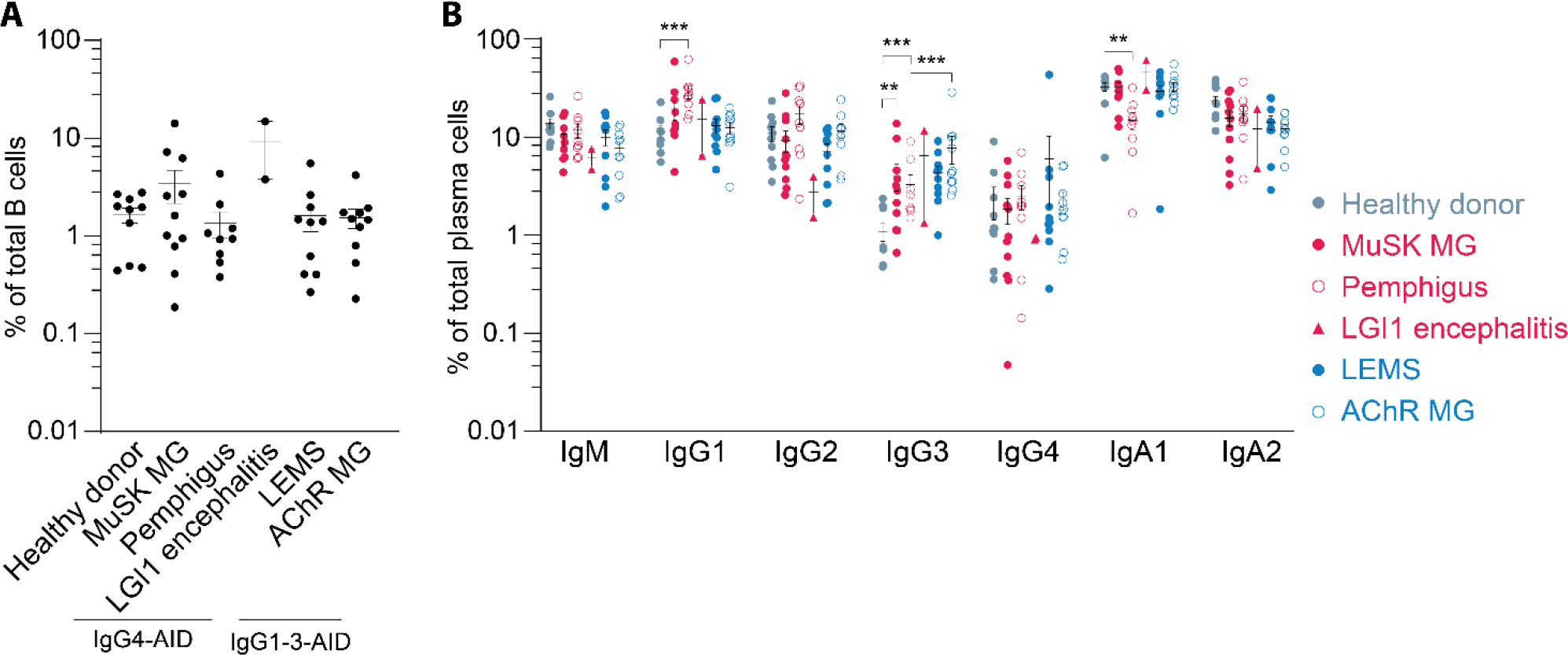
Plasma cell fractions of IgG4-AID and IgG1-3-AID patients. **(A)** Overall plasma cell (CD19^+^CD20^+/-^CD138^+/-^) counts normalized to total B cell counts. **(B)** Plasma cell counts normalized to total plasma cell counts stratified per B cell receptor IgH (sub)class. Groups were compared using one-way ANOVA on log-transformed data. If significant, Student’s t-tests were performed to compare IgG4-AID groups with other patient groups and all patient groups with healthy controls. For all panels, IgG4-AIDs were compared to both healthy controls and IgG1-3-AIDs (* p<0.05; ** p<0.01; *** p<0.005).

**Figure 4.**
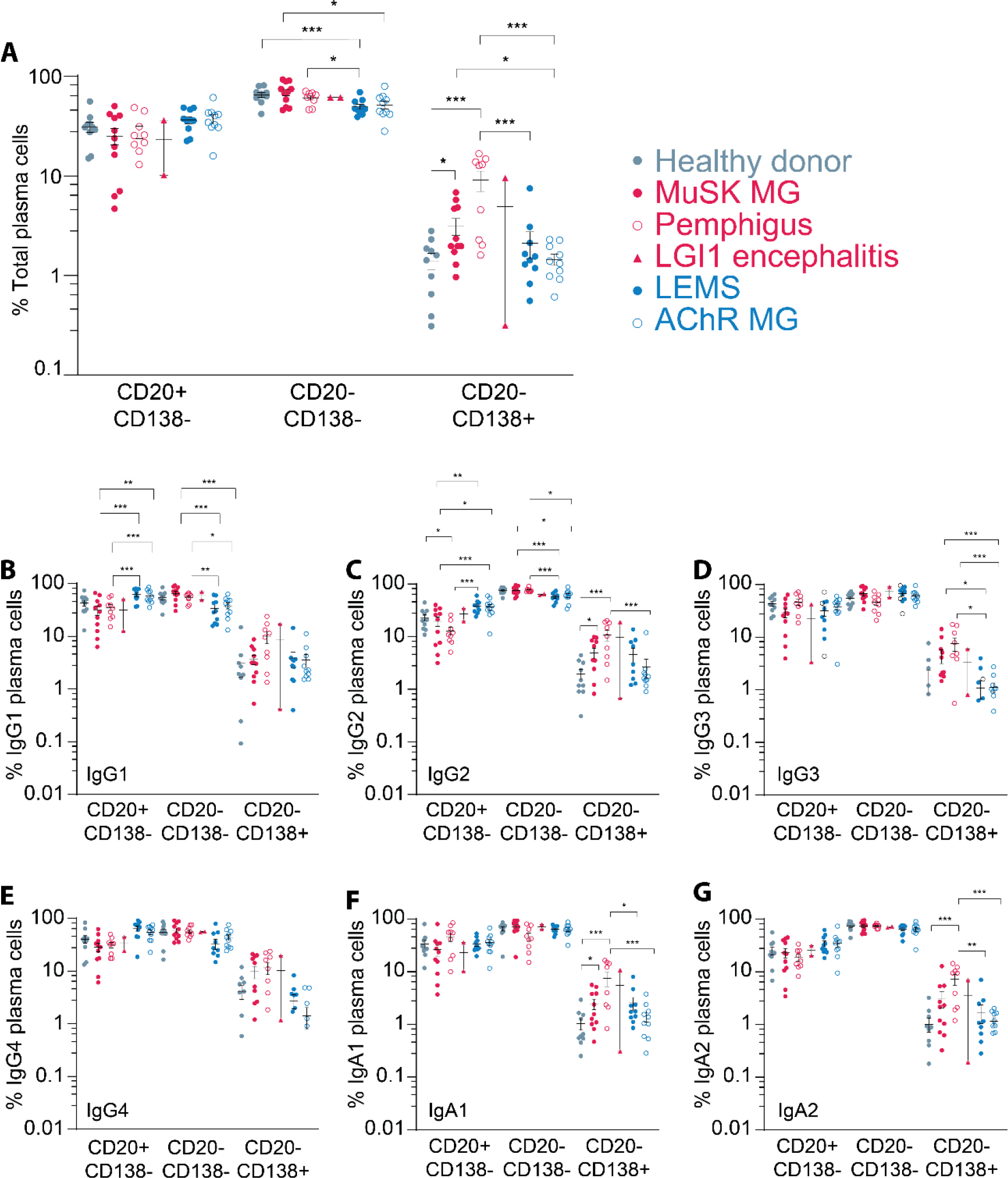
Distribution of plasma cell maturation stages of IgG4-AID and IgG1-3-AIDs. **(A)** Overall plasma cell (CD19^+^CD20^+/-^CD138^+/-^) counts normalized to total B cell counts and subdivided for maturation status normalized to total plasma cell counts. The same plasma cell populations are shown for IgG1 **(B)**, IgG2 **(C)**, IgG3 **(D)**, IgG4 **(E)**, IgA1 **(F)** and IgA2 **(G)**. For all panels, IgG4-AIDs were compared to both healthy controls and IgG1-3-AIDs by one-way ANOVA followed by unpaired Student’s t-test (* p<0.05; ** p<0.01; *** p<0.005).

Mature plasma cells of IgG4-AID patients seem to segregate in two populations. Immunosuppresive treatment may alter B cell compartment composition. The acute nature of these autoimmune disease often requires patients to start immunomodulatory treatment quickly after diagnosis. The samples included in this study were prioritized on no or low amounts of immunosuppressive treatment. However, some patients did receive prednisone, rituximab or azathioprine (Sup. Table 1). To investigate if these treatments biased our analysis we plotted the data including the treatment (Sup. Fig. 4-5). The low numbers in each treatment category prevent statistical analysis, but this plot may suggest that untreated patients have higher fractions of mature plasma cells and that treatment may lowered their numbers.

**Figure 5.**
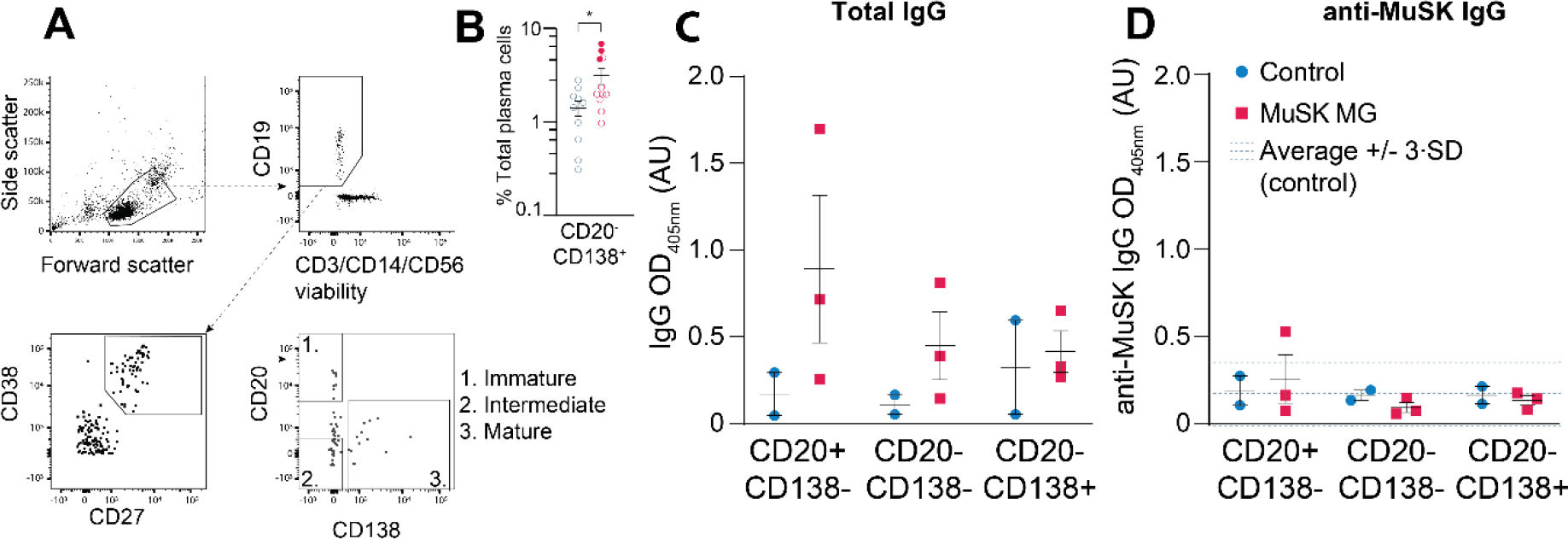
Plasma cell subset sorting and follow-up culture to investigate autoreactivity. **(A)** Representative flow plots and gating strategy of plasma cell sorting into CD20^+^CD138^-^, CD20^-^CD138^-^ and CD20^-^CD138^+^ subsets. **(B)** Excerpt from Fig. 4A, patients selected for plasma cell sort are shown as filled dots. **(C)** Total IgG titers produced by plasma cell subsets after culturing for 14 days. **(D)** MuSK-reactive IgG titers produced by plasma cell subsets after culturing for 14 days. The dashed lines represent the average anti-MuSK IgG titer of the control sample with a range of 3 standard deviations.

### Autoreactivity is not enriched in any of the circulating plasma cell maturation stages

To investigate whether these CD20^-^CD138^+^ plasma cells of IgG4-AID patients include autoantibody-producing cells, we sorted CD20^+^CD138^-^, CD20^-^CD138^-^ and CD20^-^CD138^+^ plasma cells of MuSK MG patients and compared them to healthy controls (Fig. 5A). We selected 3 untreated MuSK MG patients with relatively high numbers of mature plasma cells (marked in Fig. 5B and Sup. Table 1) for this experiment. After sorting, these populations were taken into culture to collect supernatants for screening on MuSK-specific antibodies (18). Despite detecting total IgG in supernatants of all three plasma cell subsets (Fig. 5C), no MuSK-specific IgG was found in any of the subsets except for 1 patient in the CD20^+^CD138^-^ population (Fig. 5D).

## 3. Discussion

To investigate whether predominance of IgG4 autoantibodies in IgG4-AIDs is caused by aberrations in B cell development or class switching, we compared the full peripheral blood B cell compartment of the IgG4-AIDs MuSK MG, pemphigus, LGI1 and CASPR2 encephalitis with the IgG1-3-AIDs AChR MG and LEMS, as well as healthy controls. Generally, B cell relative frequencies, and therefore also B cell development, were normal across all autoimmune diseases tested. B cell numbers from our healthy donors matched well with previous reports (49). IgG4^+^ memory B cell or IgG4^+^ plasma cell fractions were not increased in IgG4-AID patients. This suggests that IgG4-AID patients do not have aberrant B cell receptor class switching favoring an IgG4 response. This is in line with studies showing that IgG4 serum levels are only mildly, if at all, increased in IgG4-AID patients (22–24). Generalized IgG4 B cell fractions in IgG4-AID patients being comparable to healthy controls suggests that the IgG4 predominance in autoimmune responses is selective, antigen-specific and perhaps antigen-driven (see below). The HLA class II associations as well as the favorable response to rituximab across IgG4-AIDs, in combination with the data presented here, further support the hypothesis of an overarching immunophenotype across IgG4-AIDs.

This data also strengthens the idea that IgG4-AIDs represent a different disease entity from IgG4-related diseases (IgG4-RDs). IgG4-RD are hallmarked by increased numbers of circulating IgG4^+^ memory B cells and IgG4^+^ plasmablasts coupled to high IgG4 serum titers and tissue fibrosis (50). While various autoantibodies have been found in IgG4-RD patients, these are mainly of the IgG1 subclass and are not known to correlate consistently with the disease (51–53). Serum IgG4 autoantibody titers do correlate with disease severity in IgG4-AIDs and cause disease upon passive transfer (20, 21, 54). Despite the central role of IgG4 in both disease groups, IgG4-AIDs and IgG4-RDs should not be considered part of the same disease spectrum.

IgG4-AID patients were found to have an overall increase in mature (CD20^-^CD138^+^) plasma cells, but this increase was not unique to IgG4^+^ cells. Increased mature plasma cells were previously reported in some (55), but not all (56) studies on AChR MG patients. We did not observe this in AChR MG patients included in this study. Plasma cell numbers decrease with age (49, 57). The differences between the AChR MG studies may be explained by this confounding effect. We did not observe any age-dependent plasma cell decrease in our study population (Sup. Fig. 6). Increased CD138^+^ plasma cells are a hallmark of several chronic AIDs (58, 59) and both MuSK MG and pemphigus likely fit the same classification. Why the other AIDs did not show increased mature plasma cell numbers is not known.

CD138^+^ plasma cells are usually contained within the bone marrow and are considered responsible for long term immunity (60). This fully matured plasma cell subset shows the highest levels of antibody secretion. Why certain immune responses induce such mature responses is not fully understood, but has been reported at day 7 post-vaccination for common vaccines against pathogens such as mumps, tetanus, pertussis and measles (45, 60–62). This suggests an immune response timing-dependent migration. In IgG4-AIDs these mature plasma cell levels may be increased as a result of: 1) a net increase in their numbers due to a change in antigen-independent maturation, 2) stronger migratory signals that may stimulate these cells to leave the bone marrow and become increased in PBMCs, possibly during migration towards sites with high target antigen availability, or, 3) these may contain the autoantibody producing plasma cell subsets which increase their overall numbers due to chronic activation. Although there was no selective increase of IgG4 subclass mature plasma cells, we tested the third hypothesis by sorting the early, intermediate and mature plasma cell populations of MuSK MG patients and testing for autoantibody production. We did not find evidence for autoreactivity in any of these subsets. Although we detected robust total IgG production in these cultures, we cannot exclude a technical limitation as sorting and subsequently culturing these delicate plasma cell populations is challenging (63). A role for these mature plasma cells in the autoimmune response may be considered unlikely due to the fact that they do not express CD20, but rituximab (anti-CD20 therapy) treatment is usually effective in IgG4-AIDs (25–28). IgG4 responses are mostly mediated by short-lived plasma cells which do not migrate to the bone marrow (64). Bone marrow holds many of the cell types that contribute to humoral immunity. PBMCs may therefore not accurately reflect aberrations present in the bone marrow compartment. Whether peripheral or bone marrow mature plasma cells play a role in the pathophysiology of IgG4-AIDs requires further investigation.

Pre-germinal center subsets of immature and naïve CD5^+^ B cells were decreased in IgG4-AIDs. Previous work has shown that immunosuppressants, especially azathioprine, selectively lower naïve CD5^+^ B cell counts, which may explain this observation (39). Indeed, azathioprine treated patients show the most severe decrease in naïve CD5^+^ B cell numbers in our cohorts (Figure 1). Other immunomodulatory treatments may also significantly bias immunophenotyping analysis. The severity of the IgG4-AIDs requiring rapid treatment combined with a poor response to symptomatic treatments made inclusion of untreated patients challenging (65, 66). We therefore aimed to include as much immunosuppressive treatment naïve patients for this study, but, due to the rarity of these samples, were compelled to also include some who received (low amounts of) immunosuppression. Although the included patients all experienced significant disease symptoms, we cannot fully exclude that in some patients the treatment regimens affected the B cell compartment. Future immune profiling studies should aim to only include untreated individuals whenever possible.

In the memory B cell compartment, we found no differences between any of the patient groups in both total cell abundance or when stratified for the switched (IgM^-^/D^-^CD27^+^) or double negative (IgM^-^ /D^-^CD27^-^) subsets. We did observe a significant decrease of IgG4^+^ atypical (IgM^-^/D^-^CD21^-^CD27^-^) memory B cells in MuSK MG patients when compared to healthy controls or IgG1-AID patients. Like IgG4 responses in general, atypical B cell counts increase with prolonged and repeated antigen exposure (67) and are associated with several autoimmune diseases (68–71). However, we did not detect this in our study for the other AIDs.

The question thus remains why these autoimmune diseases are characterized by predominant IgG4 autoantibody responses. One possibility is that the antigen itself directs the response towards IgG4. Certain antigens are known to drive IgG4 responses, such as bee venom and certain biologicals (72). Specifically for pemphigus there is evidence of desmoglein 1/3 autoantibody development following exposure to walnut or sand fly antigens (36–38). There is no evidence yet for comparable molecular mimicry events in other IgG4-AIDs. However, given the strong correlation between these antigens and IgG4, molecular mimicry could be a plausible factor in the etiology of other IgG4-AIDs.

## 4. Methods

### 4.1 Study population

AID patients with Muscle-specific kinase (MuSK) myasthenia gravis (MG), acetylcholine receptor (AChR) MG, Lambert-Eaton myasthenic syndrome (LEMS) or pemphigus (vulgaris, foliaceus and paraneoplastica) were recruited from the Leiden University Medical Centre (2017–2021). We obtained blood samples for 10 patients per disease except for MuSK MG, for which we obtained 11 samples. Two Contactin-associated protein-like 2 (CASPR2) encephalitis and three leucine-rich glioma inactivated (LGI1) encephalitis patients were recruited from the Erasmus University Medical Center (2019–2021). Patients were included based on the presence of symptoms matching MG, pemphigus, encephalitis or LE (73–75) and a positive titer on a serological test for the respective autoantibody upon standard clinical testing. Patients were excluded if they had received rituximab treatment within the past 12 months. Blood samples were also obtained from 10 age- and sex-matched healthy controls. These healthy controls were recruited by the LUMC Voluntary Donor Service (LuVDS). Figure 1 and Supplemental Table 1 provide a detailed overview of the study population.

### 4.2 Isolation of peripheral blood mononuclear cells

Peripheral blood mononuclear cells (PBMCs) were isolated from 60-90 ml of sodium-Heparin anticoagulated peripheral blood samples by Ficoll-amidotrizoate density gradient centrifugation. Following isolation, cells were immediately frozen at -80° C at a density of 5-10·10^6^ per ml in Recovery Cell Freezing medium (Thermo Fisher Scientific, Waltham, MA, USA) in a Mr. Frosty Freezing Container (Thermo Fisher Scientific, Waltham, MA, USA) for 24 hours before transfer to liquid nitrogen storage.

### 4.3 Flow cytometry

The B cell subsets and B cell receptor (sub)classes were identified in freshly thawed PBMCs using the standardized EuroFlow 12-color IgH-isotype B-cell tube (49, 76, 77), with the exception of CD62L which was replaced by a Zombie Yellow cell viability stain (BioLegend, San Diego, CA, USA) (see Sup. Table 2 for a detailed overview). At least 10·10^6^ cells were stained for 30 minutes in the dark in 100 µl (75 μl EuroFlow B cell tube mix and 25 μl Cytognos isotype mix) staining solution according to the EuroFlow SOP for sample preparation and staining of markers followed by immediate analysis (www.EuroFlow.org).

Flow cytometry was performed on a BD FACS LSR Fortessa 4L (BD Biosciences, San Jose, CA, USA) at the Flow cytometry Core Facility (FCF) of Leiden University Medical Center (LUMC) in Leiden, Netherlands (https://www.lumc.nl/research/facilities/fcf). At least 3·10^6^ cells were acquired per sample. Instrument setup was according to the EuroFlow standardized operating procedures (78). See Sup. Fig. 1 for a detailed overview of the gating strategy.

Plasma cells were sorted on a BD FACSAria 3 cell sorter (BD Biosciences, San Jose, CA, USA). The gating strategy is detailed in Sup Fig. 2. In brief, freshly thawed PBMCs were stained for expression of CD38, CD20, CD138 and CD19. Viable cells were identified using Zombie Green cell viability stain (BioLegend, San Diego, CA, USA). See Sup. Table 3 for a detailed overview of the staining. A dump channel for T cells, NK cells and macrophages was created by staining PBMCs for expression of CD3, CD14 and CD56.

### 4.4 Plasma cell culture

After bulk sorting, plasma cell subtypes were cultured in RPMI1640 (Thermo Fisher Scientific, Waltham, MA, USA) supplemented 10% heat-inactivated fetal bovine serum, IL-6 (10 ng/ml; Thermo Fisher Scientific, Waltham, MA, USA), IL-21 (50 ng/ml; Thermo Fisher Scientific, Waltham, MA, USA), IFN-α (100 U/ml; Merck, Rahway, NJ, USA), BAFF (20 ng/ml; Miltenyi Biotec, Bergisch Gladbach, NRW, Germany), chemically defined lipid mixture 1 (1/200; Thermo Fisher Scientific, Waltham, MA, USA), MEM amino acid solution (1×; Sigma-Aldrich, St. Louis, MO, USA) on γ-irradiated M2-10B4 stromal cells (1.5*10^4^/well, 100 μl/well) in 96-well plates (79). Every 7 days, 50 μl culture supernatant was aspirated for IgG detection (anti-MuSK and total IgG) and replaced with fresh medium.

### 4.5 ELISA

Plasma cell culture medium samples were screened for antibody production using a total human IgG ELISA assay and for MuSK reactivity using a previously described MuSK ELISA assay (80).

### 4.6 Statistics

Flow cytometry data was blinded and then analyzed using Infinicyt 2.0 (Cytognos, Salamanca, Spain). Statistical analyses were performed using Prism 9 (GraphPad Software, San Diego, CA, USA). Data was log transformed and significance was assessed by one-way ANOVA followed by unpaired Student’s t-test unless otherwise specified. IgG4-AID patients were compared to healthy controls and to IgG1-3-AID patients. P-values below 0.05 are considered statistically significant (* p<0.05; ** p<0.01; *** p<0.005).

### 4.7 Study approval

This study was approved by the local medical ethics committee of the Leiden University Medical Centre (CME protocolnumber P17.011). All subjects provided written informed consent prior to participation and experiments were in accordance with the Declaration of Helsinki, including current revisions, and Good Clinical Practice guidelines.

### 4.7 Data availability

All raw flow cytometry data for this work is directly accessible via the Flow Repository under experiment ID FR-FCM-Z6K2 (81). Derived data supporting the findings of this study are available from the corresponding author on request.

## 5. Author contributions

**Laurent M. Paardekooper:** Formal Analysis, Writing – Original Draft, Visualization, Data Curation

**Yvonne E. Fillié-Grijpma:** Investigation, Formal Analysis, Data Curation

**Alita J. van der Sluijs-Gelling:** Investigation, Formal Analysis

**Mihaela Zlei:** Methodology, resources

**Remco van Doorn:** Resources

**Maarten H. Vermeer:** Resources

**Manuela Paunovic:** Resources

**Maarten J. Titulaer:** Resources

**Silvère M. van der Maarel:** Writing – Review & Editing

**Jacques J.M. van Dongen:** Conceptualization, Writing – Review & Editing, Resources, Methodology

**Jan J. Verschuuren:** Conceptualization, Writing – Review & Editing, Resources

**Maartje G. Huijbers:** Conceptualization, Writing – Review & Editing, Funding Acquisition, Supervision, Project Administration

## Acknowledgements

The authors gratefully acknowledge dr. Jelle Goeman for his advice on statistical procedures.

The authors gratefully acknowledge the Flow cytometry Core Facility (FCF) of Leiden University Medical Center (LUMC) in Leiden, the Netherlands (https://www.lumc.nl/research/facilities/fcf), coordinated by dr. K. Schepers and M. Hameetman, run by the FCF Operators E.F.E de Haas, J.P. Jansen, D.M. Lowie, S. van de Pas, and G.IJ. Reyneveld (Directors: Prof. F.J.T. Staal and Prof. J.J.M. van Dongen) for technical support.

The authors gratefully acknowledge the LUMC Vrijwillige Donoren Service (LuVDS) for providing healthy donor material.

The authors acknowledge the support of patient partners, private partners and active colleagues of the T2B consortium; see website: www.target-to-b.nl.

The collaboration project is financed by the PPP Allowance made available by Top Sector Life Sciences & Health to Samenwerkende Gezondheidsfondsen (SGF) under project code LSHM18055-SGF to stimulate public-private partnerships and co-financing by health foundations that are part of the SGF.

## Conflict of interest

JV, SM, MH are coinventors on MuSK-related patents. LUMC and JV, SM and MH receive royalties from these patents. SM is a Board member for Renogenyx. LUMC receives royalties on a MuSK ELISA. JV is consultant for Argenx, Alexion, NMD Pharma. MH receives financial support from the LUMC (OIO 2017, Gisela Their Fellowship 2021), Top Sector Life Sciences & Health to Samenwerkende Gezondheidsfondsen (LSHM19130), Prinses Beatrix Spierfonds (W.OR-19.13). MJT is member of the European Reference Network for Rare Immunodeficiency, Autoinflammatory and Autoimmune Diseases–Project ID No 739543 (ERN-RITA; HCP Erasmus MC). MJT has filed a patent, on behalf of the Erasmus MC, for methods for typing neurological disorders and cancer, and devices for use therein, and has received research funds for serving on a scientific advisory board of Horizon Therapeutics, for consultation at Guidepoint Global LLC, for consultation at UCB, for teaching colleagues at Novartis. MJT has received an unrestricted research grant from Euroimmun AG, and from CSL Behring. JJMvD reports to be the chairman of the EuroFlow Consortium. JJMvD is also listed as inventor on the patent “Means and methods for multiparameter cytometry-based leukocyte subsetting” (PCT/NL2020/050688, filing date 5 November 2019), owned by the EuroFlow scientific consortium; JJMvD reports an Educational Services Agreement from BD Biosciences (San José, CA) and a Scientific Advisor Agreement with Cytognos/BD Biosciences; all related fees and honoraria are for the involved university departments at Leiden University Medical Center and University of Salamanca. The remaining authors declare no interests. The LUMC is part of the European Reference Network for Rare Neuromuscular Diseases [ERN EURO-NMD] and the Netherlands Neuromuscular Center.

The other authors have declared that no conflict of interest exists.

