## Supplemental Figures for "Autoantibody subclass predominance is not driven by aberrant class switching or impaired B cell development"

### 8. Supplemental data

**Sup. Table 1. Study population (individual subjects)**

Disease severity for both MuSK and AChR MG patients was assessed using the Quantitative Myasthenia Gravis (QMG) (82) and Myasthenia Gravis Activities of Daily Living Scale (MG-ADL) (83) scoring methods.

| | Age at blood draw (years) | Sex | Disease duration (months) | Severity | Serology (antigen, U/ml) | Treatment at blood draw | Remarks |
|---|---|---|---|---|---|---|---|
| **MuSK MG** | | | | **QMG; MG-ADL** | | | |
| PCMG005 (sorted) | 79 | M | 84 | 6; 3 | Anti-MuSK (0,61) | unknown | QMG and MG-ADL 2 months prior to blood draw |
| PCMG008 | 62 | M | 0 | 2; 0 | Anti-MuSK (>0,7) | Prednisone, plasmaferesis (5 days) | |
| PCMG009 | 69 | F | 240 | n.a. | Anti-MuSK (>0,7) | Prednisone. Azathioprine before. | |
| PCMG018 (sorted) | unknown | F | unknown | unknown | unknown | unknown | No patient data available at LUMC |
| PCMG020 | 31 | F | 60 | n.a. | Anti-MuSK (>0,7) | Prednisone, cellcept, rituximab. Azathioprine before. | |
| PCMG021 | 69 | M | 156 | 7; 10 | Anti-MuSK (>0,7) | Cellcept, rituximab | |
| PCMG025 | 32 | F | 240 | n.a. | Anti-MuSK (11,7) | Cellcept. Rituximab before. | |
| PCMG030 | 56 | M | 8 | 5; 4 | Anti-MuSK (0,69) | Prednisone, azathioprine, IVIG (5 days) | |
| PCMG033 | 74 | F | 5 | 9; 3 | Anti-MuSK (2,2) | Prednisone, azathioprine | |
| PCMG036 (sorted) | 27 | F | 48 | n.a. | Anti-MuSK (0,9) | Untreated | |
| PCMG037 | 43 | F | 10 | 5; 6 | Anti-MuSK (>12,0) | Untreated | |
| **Pemphigus** | | | | **1-3 (mild-severe)** | | | |
| PVBcel001 | 66 | F | 8 | 1 | Unknown | Prednisone 50mg | pemphigus vulgaris |
| PVBcel002 | 75 | F | 4 | 2 | Dsg1 (97), Dsg3 (35) | Prednisone 80mg | paraneoplastic pemphigus |
| PVBcel003 | 54 | M | 5 | 1 | Dsg1 (44) | Rituximab | pemphigus foliaceus |
| PVBcel004 | 69 | M | 5 | 2 | Dsg1 (71) | Prednisone 30 mg | pemphigus foliaceus |
| PVBcel005 | 29 | M | 29 | 3 | Dsg1 (>150), Dsg3 (>150) | Clobetasol lotion. Rituximab >1 year before. | pemphigus vulgaris |
| PVBcel006 | 52 | M | 40 | 2 | Dsg3 (125) | Prednisone 50mg | pemphigus vulgaris |
| PVBcel007 | 26 | M | 2 | 3 | Dsg1 (117), Dsg3 (>150) | Prednisone 30mg | pemphigus vulgaris |
| PVBcel008 | 50 | F | 7 | 3 | Dsg1 (>15), Dsg3 (>150) and BP180 | Prednisone 50mg | pemphigus vulgaris and bulleus pemphigoid |
| PVBcel009 | 62 | M | 13 | 2 | Dsg1 (5), Dsg3 (114) | Prednisone 5mg | pemphigus vulgaris |
| PVBcel010 | 62 | F | 4 | 2 | Dsg1 (>100) | Prednisone 30mg | pemphigus foliaceus |
| **LGI1 limbic encephalitis** | | | | **1-3 (mild-severe)** | | | |
| PBMC060721_LGI1 | 62 | M | 3 | 2 | LGI1 (++; VGKC titer 626) | Untreated | |
| Sample210920_LGI1 | 48 | F | 7 | 2 | LGI1 (+; VGKC titer 277) | IVIg course started 21 days before (day -21 to -17), just before second course | |
| **CASPR2 neuromyotonia** | | | | **1-3 (mild-severe)** | | | |
| Sample080120_CASPR2 | 65 | M | 10 | 2 | CASPR2 (++; VGKC 301) | Azathioprine 2d 75mg; steroids tapered down >3 months before. | Morvan syndrome; relapse at time of blood draw |
| Sample110121_CASPR2 | 57 | M | 1 | 2 | CASPR2 (+; VGKC <50) | Untreated; ivMP, PLEX and thymoma resection >1 yr before (for AChR MG) | Morvan syndrome |
| Sample161219_CASPR2 | 69 | M | 36 | 2 | CASPR2 (++; VGKC 412) | Azathioprine; ivMP and IVIg 1 yr before; oral steroids tapered down >6mo before | Relapse at time of blood draw |
| **LEMS** | | | | | | | |
| PCMG011 | 56 | M | 288 | n.a. | VGCC (359) | Hydrocortisone (on unrelated indication) | |
| PCMG012 | 55 | M | 12 | n.a. | VGCC (151) | Untreated | |
| PCMG014 | 74 | F | 24 | n.a. | VGCC (286) | Untreated | |
| PCMG015 | 65 | M | 120 | n.a. | VGCC (207) | Prednison | |
| PCMG017 | 52 | F | 60 | n.a. | VGCC (179) | Untreated | |
| PCMG032 | 56 | F | 84 | n.a. | VGCC (134) | Untreated | |
| PCMG038 | 65 | F | 180 | n.a. | VGCC (99) | Untreated. methotrexate, prednisone and IVIG before. | |
| PCMG043 | 59 | F | 36 | n.a. | VGCC (67) | Azathioprine, IVIG | |
| PCMG049 | 55 | F | 36 | n.a. | VGCC (317) | Untreated | |
| PCMG050 | 49 | F | 96 | n.a. | VGCC (299) | Untreated | |
| **AChR MG** | | | | **QMG; MG-ADL** | | | |
| PCMG013 | 56 | F | 60 | unknown | AChR (>5.0) | Untreated | No patient data available at LUMC |
| PCMG022 | 69 | M | 132 | n.a. | AChR (5.5) | Prednison | |
| PCMG023 | 72 | M | 12 | n.a. | AChR (>5.5) | Prednison | |
| PCMG024 | 36 | F | 156 | n.a. | AChR (>5.5) | Untreated | |
| PCMG027 | 64 | M | 12 | 8; 8 | AChR (>5.5) | Untreated | |
| PCMG028 | 66 | F | 132 | n.a. | AChR (5.5) | Untreated. Prednisone before. | |
| PCMG042 | 18 | F | 24 | 31; 16 | AChR (>5.5) | Prednisone, IVIG | |
| PCMG044 | 62 | F | 12 | 9; 4 | AChR (5.3) | Prednisone, IVIG | |
| PCMG045 | 40 | F | 48 | 25; 11 | AChR (>5.5) | Untreated | |
| PCMG046 | 79 | F | 4 | 19; 14 | AChR (>5.5) | Prednisone | |
| **Healthy control** | | | | | | | |
| LuVDS0001 | 68 | M | - | - | - | - | |
| LuVDS0018 | 46 | M | - | - | - | - | |
| LuVDS0022 | 66 | F | - | - | - | - | |
| LuVDS0028 | 57 | M | - | - | - | - | |
| LuVDS0030 | 54 | F | - | - | - | - | |
| LuVDS0049 | 62 | F | - | - | - | - | |

**Sup. Table 2. EuroFlow B cell panel**

| Marker | Fluorochrome | Clone | Source | Cat. No. | µl/test (100 µl) |
|---|---|---|---|---|---|
| CD5 | PE-Cy7 | L17F12 | BD Biosciences | 348810 | 6 |
| CD19 | BV786 | SJ25C1 | BD Biosciences | 563325 | 4 |
| CD20 | PE-CF594 | 2H7 | BD Biosciences | 562550 | 5 |
| CD21 | BV711 | B-ly4 | BD Biosciences | 563163 | 5 |
| CD24 | BV650 | ML5 | BD Biosciences | 563720 | 5 |
| CD27 | BV421 | M-T271 | BD Biosciences | 562513 | 2 |
| CD38 | APC-H7 | HB-7 | BD Biosciences | 656646 | 3 |
| CD45 | AF700 | HI30 | BD Biosciences | 560566 | 10 |
| CD138 | PE-Cy7 | MI15 | BioLegend | 356513 | 5 |
| SmIgD | FITC | IA6-2 | BioLegend | 348205 | 1,25 |
| SmIgD | APC | IA6-2 | BD Biosciences | 561303 | 4 |
| SmIgM | BV510 | MHM-88 | BioLegend | 314521 | 2 |
| SmIgA1 | APC | SAA1 | Cytognos | CYT-IGS1 | 25 of mix |
| SmIgA1 | PerCP-Cy5 | SAA1 | Cytognos | CYT-IGS1 | |
| SmIgA2 | PerCP-Cy5 | SAA2 | Cytognos | CYT-IGS1 | |
| SmIgG1 | PE | SAG1 | Cytognos | CYT-IGS1 | |
| SmIgG2 | FITC | SAG2 | Cytognos | CYT-IGS1 | |
| SmIgG2 | PE | SAG2 | Cytognos | CYT-IGS1 | |
| SmIgG3 | FITC | SAG3 | Cytognos | CYT-IGS1 | |
| SmIgG4 | APC | SAG4 | Cytognos | CYT-IGS1 | |
| Viability | Zombie Yellow | - | BioLegend | 423103 | 0.2 |

By: Blanco et al., 2018; Blanco et al., 2019; van Dongen et al., 2019

**Sup. Table 3. Plasma cell sort panel**

| **Marker** | **Fluorochrome** | **Clone** | **Source** | **Cat. No.** | **µl/test (600 µl)** |
|---|---|---|---|---|---|
| CD19 | BV421 | SJ25C1 | BioLegend | 363017 | 6 |
| CD20 | AF700 | 2H7 | BD Biosciences | 560613 | 12 |
| CD27 | APC-H7 | M-T271 | BD Biosciences | 560223 | 24 |
| CD38 | APC | HB-7 | BioLegend | 356605 | 6 |
| CD138 | PE | MI15 | BioLegend | 356503 | 6 |
| CD3 | FITC | UCHT1 | BD Biosciences | 561806 | 6 |
| CD14 | FITC | M5E2 | BD Biosciences | 561712 | 6 |
| CD56 | FITC | HCD56 | BioLegend | 318304 | 6 |
| Viability | Zombie Green | - | BioLegend | 423111 | 0.4 |

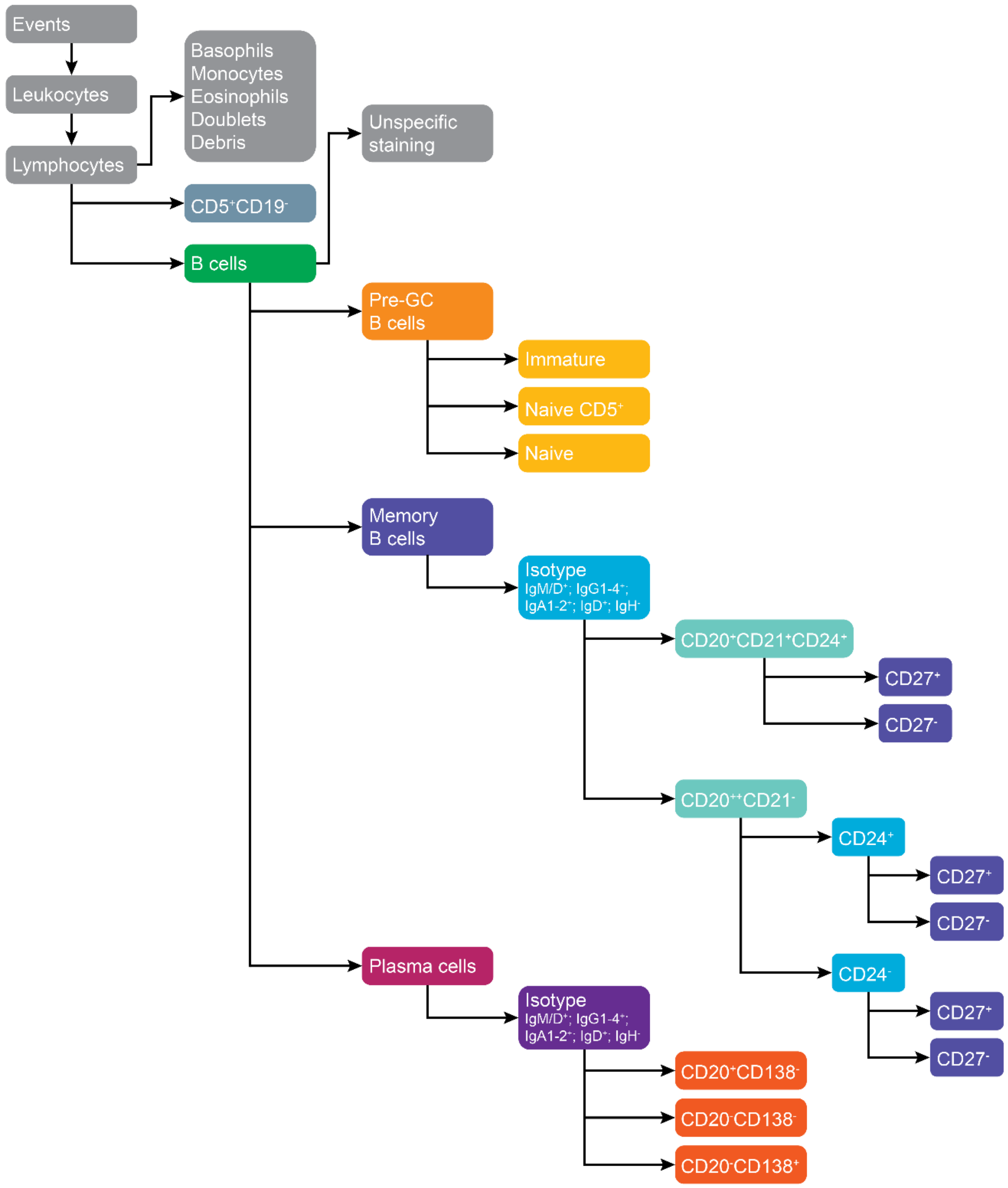


**Sup. Figure 1. Flow cytometry gating strategy**

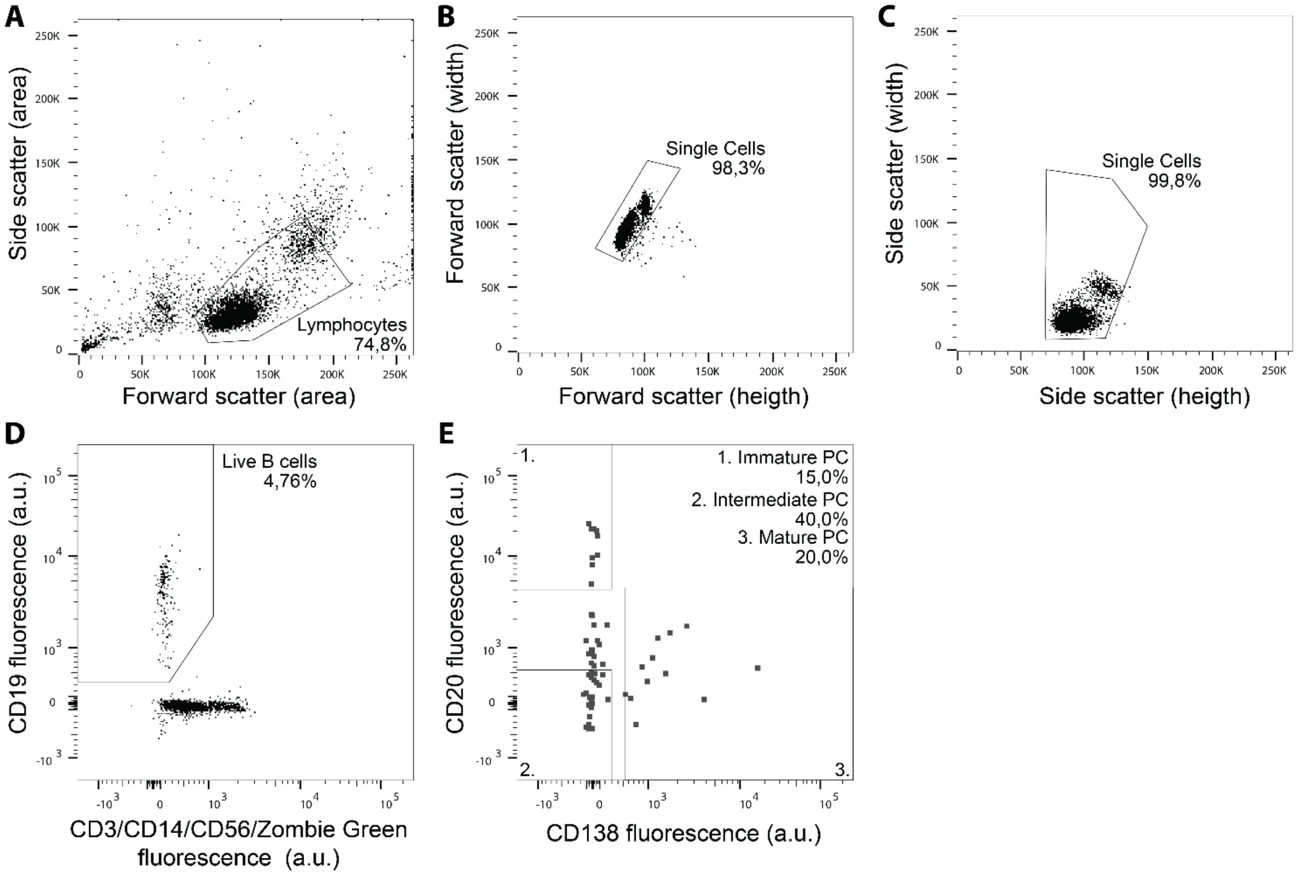


**Sup. Figure 2. Plasma cell sort gating strategy. (A)** Rough gating for lymphocytes based on forward and side scatter, **(B, C)** followed by doublet exclusion. **(D)** B cells are then gated based on $CD19^{+}CD3^{-}CD14^{-}CD56^{-}$ expression profile and negative Zombie Green viability staining. **(E)** Selection of immature ($CD20^{+}CD138^{-}$), intermediate ($CD20^{-}CD138^{-}$) and mature ($CD20^{-}CD138^{+}$) plasma cells.

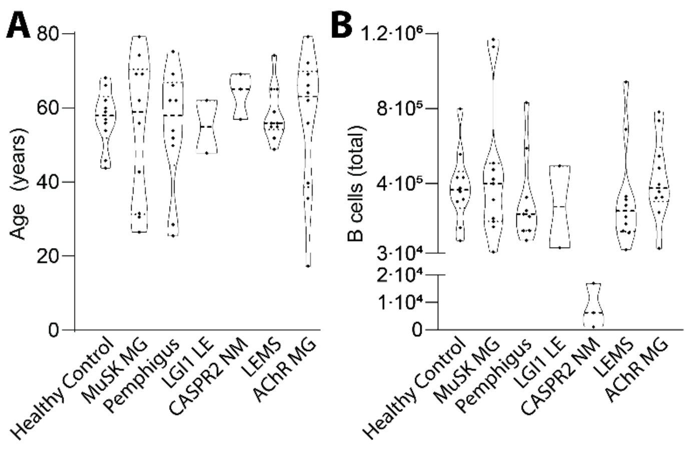

**Sup. Figure 3. Study population.** Distribution of **(A)** age and **(B)** total number of B cells analyzed for each donor subcategory. Dotted lines represent median and interquartile ranges. Point distribution width is proportionate to the number of points at each Y value. No significant differences were found (one-way ANOVA; p=0.963, p=0.263, respectively)

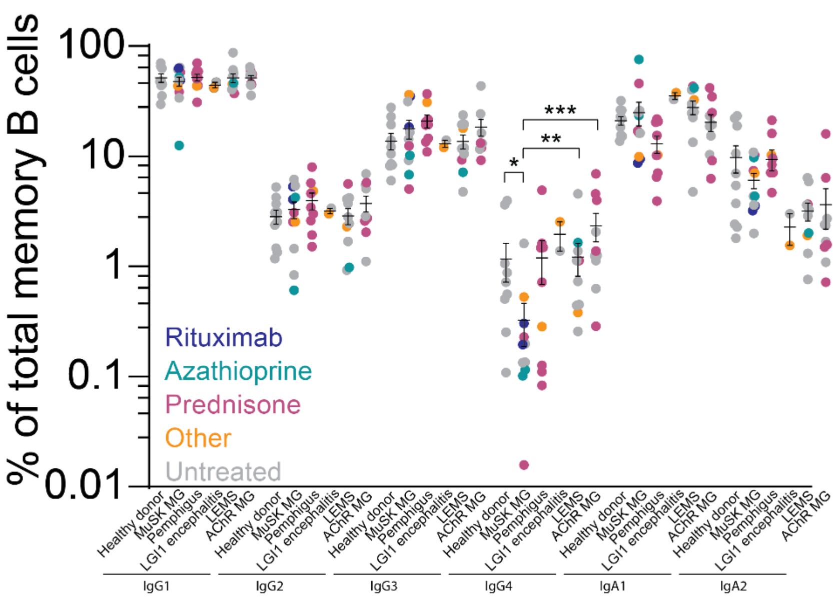


**Sup. Figure 4. Atypical memory B cell fractions with treatment status**

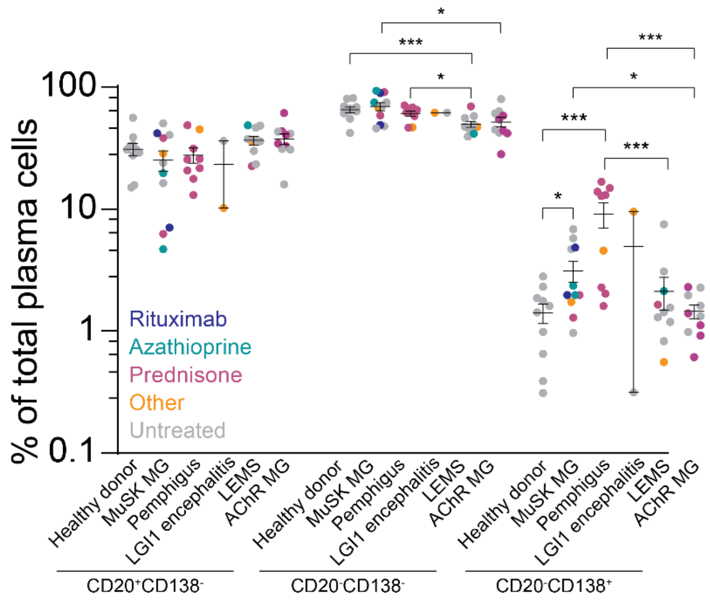


**Sup. Figure 5. Plasma cell maturation stages of AID patients with treatment status.**

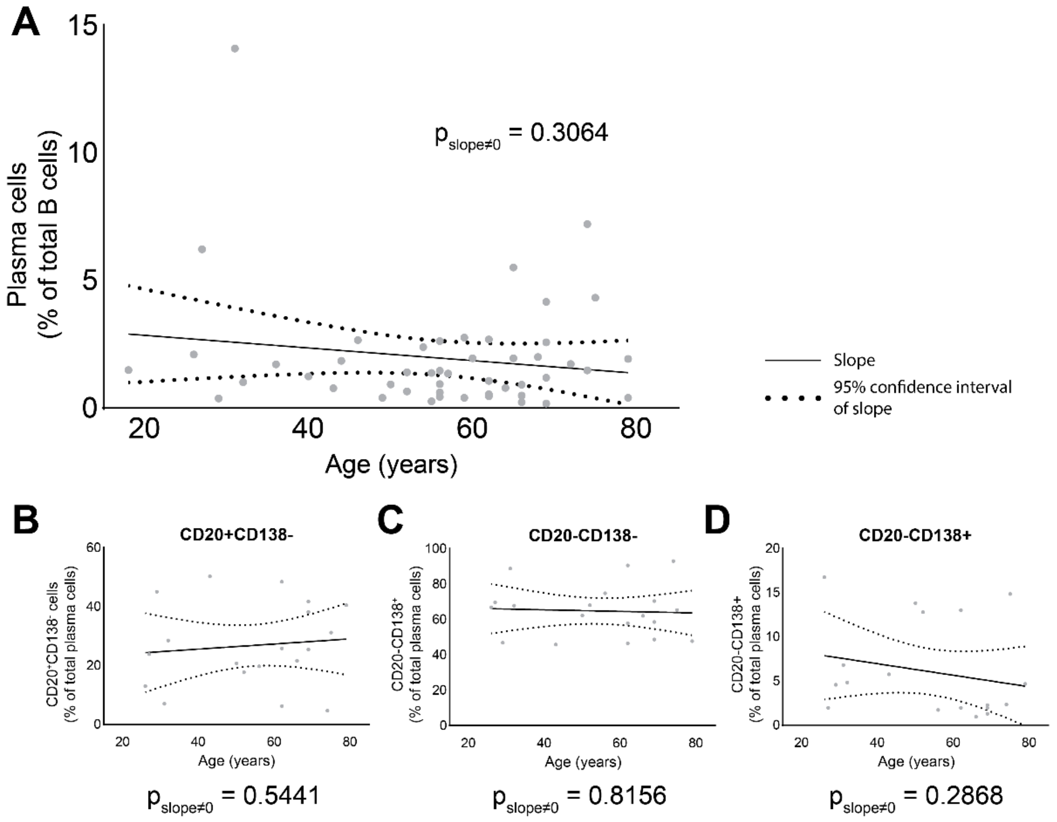


**Sup. Figure 6. Linear regression analysis of plasma cell percentage versus age. (A)** Simple linear regression on plasma cell percentage of total B cells for each donor versus donor age in years. Dotted lines represent the 95% confidence interval of the slope. Deviation from zero of the slope was found non-significant ($p = 0.3064$). **(B-D)** As A but for $CD20^+CD138^-$, $CD20^-CD138^-$ and $CD20^-CD138^+$ subsets, respectively. Deviation from zero of the slop was found non-significant in all subsets ($p = 0.5441$; $p = 0.8156$ and $p = 0.2868$, respectively).
